# Your pleasure is mine; when people share a musical emotional experience during a live music performance in a concert hall

**DOI:** 10.1101/2021.03.26.436975

**Authors:** Thibault Chabin, Damien Gabriel, Alexandre Comte, Emmanuel Haffen, Thierry Moulin, Lionel Pazart

**Affiliations:** Centre Hospitalier Universitaire de Besançon, Centre d’Investigation Clinique INSERM CIC 1431, Besançon, France; Plateforme de Neuroimagerie Fonctionnelle et Neurostimulation Neuraxess, Centre Hospitalier Universitaire de Besançon, Université de Bourgogne Franche-Comté, France; Laboratoire de Recherches Intégratives en Neurosciences et Psychologie Cognitive, Université Bourgogne Franche-Comté, F-25000, Besançon, France

**Keywords:** Emotional contagion, Emotional resonance, musical pleasure, musical reward, EEG hyperscanning, physiological coupling, live musical performance

## Abstract

Over the years, several publications have proposed that musical sound could be an ancestral emotional way of communication, thus positing an ancestral biological function for music. Understanding how musical emotions, and the pleasure derived from music regardless of the musical valence, can be shared between individuals is a fascinating question, and investigating it can shed light on the function of musical reward. Is the pleasure felt at the individual level transmitted on a collective level? And if so, how? We investigated these questions in a natural setting during an international competition for orchestra conductors. Participants (n=15) used a dedicated smartphone app to report their subjective emotional experiences in real time during a concert. We recorded participant’s electrodermal activity (EDA) and cerebral activity with electroencephalography (EEG). The overall behavioral real time ratings suggest a possible social influence on the reported and felt pleasure. The physically closer the participants, the more similar their reported pleasure. We estimated the inter-individual cerebral coherence, which indicates the degree of mutual cerebral information between pairs of participants in the frequency domain. The results show that when people simultaneously reported either high or low pleasure, their cerebral activities were closer than for simultaneous neutral pleasure reports. Participants’ skin conductance levels were also more coupled when reporting higher emotional degrees simultaneously. More importantly, the participants who were physically closer had higher cerebral coherence, but only when they simultaneously reported intense pleasure. We propose that mechanisms of emotional contagion and/or emotional resonance could explain why a form of ‘emotional connecting force’ could arise between people.

## 1 Introduction

Music is one of the greatest human abilities. From birth, the human brain is musical and can be shaped by musical experiences. However, even if many of the mechanisms underlying musical processing have been explained, the ancestral function of music and the reasons why music can emotionally move us and induce an indescribable pleasure are still open to debate [25]. Musical pleasure has been associated with the recruitment of the reward system; first, a study [64] showed dopamine releases in the dorso-ventral striatum, more precisely in the caudate during the anticipation phase, and secondly in the nucleus accumbens during peak pleasure associated with chills. Second, a pharmacological study [22] used dopaminergic modulations (either blocking or enhancing dopaminergic pathway) to show that dopamine releases are not only a consequence but actually one of the causes of the pleasure associated with music. What is intriguing is that music seems to confer no benefit on a biological level [22, 64, 77]. The involvement of ancestral brain circuits, which are essential for survival and are involved in motivated behaviors (sex, food, drugs, money), in musical reward processing suggests an ancestral function for music.

Since the Darwinian period, many authors have formulated theories about the origins of music. There is an ongoing debate between non-adaptationists who claim music as a human invention, a human “pleasure technology” [60], without any ancestral biological advantages, and adaptationists who have developed theories that place music as a key mechanism for different evolutionary functions such as sexual selection, interpersonal coordination, affective communication, mood regulation and so on [19, 25, 44, 40, 1]. Musical sounds have been proposed to be an ancestral means of interpersonal emotional communication derived from “shrieking and alarm calls”, capable of producing high arousal reactions. [40, 46, 1]. Several physiological and neuro-imaging studies [11, 64, 26, 28, 29] have suggested that emotional peaks related to music, such as chills and reward-related mechanisms, might be produced when a certain emotional threshold is reached [7, 23]. In line with the Mixed Origin of Music Theory (MOM Theory) [1], Loui et al., [40] proposed that “perhaps music evolved as a direct auditory pathway toward social and emotional reward centers in the brain”, thus reconnecting the latter elements. The MOM theory also suggests that the chill reactions might have reinforced the formation of auditory memory by recruiting the reward system [1], thereby explaining the origin of music-based reward system mechanisms. Finally, Nummenmaa et al., [56] argued that “music is centrally a social rather than emotional phenomenon” and that “the social dimensions of music are the key reason why we find music enjoyable in the first place”.

One area that has received relatively little attention is how musical emotions — or more precisely musical pleasure related to reward structure activation regardless of the musical valence — are shared between people, even in the absence of direct interactions. Interindividual musical emotional sharing during indirect interactions in natural situations is a fascinating question, especially as the majority of Strong Experiences with Music [24] are reported as happening in live listening situations with other people (concerts, festivals, conferences, and so on) [37]. Research has demonstrated that musical pleasure can be directly influenced by social context, not only by musical expertise. Young children of 3 to 5 years old with a very low experience of classical music “were more emotionally responsive” when listening to music with their classmates than when listening alone [32]. A large body of research has demonstrated that the social context has a real influence on live musical experience, both on a subjective level [37, 20] and on a physiological level [70, 51, 8, 50, 3]. Inter-individual motor coordination is enhanced when listening to music collectively [71, 17]. The mere presence of other people in the same environment can enhance the synchronicity of their cardiac and respiratory activities spontaneously when they share common musical emotional experiences [3]. As far as we are aware, no studies of inter-individual cerebral coupling have placed emotional sharing at the center of their hypotheses, and no objective cerebral markers are known. So, can the pleasure felt at the individual level be transmitted on a collective level? And if so, can we objectively measure it?

We addressed this question in a natural musical setting during the International Competition for Young Conductors, held in Besançon (France). The specific natural musical settings of such an event provide a favorable framework for the study of shared musical emotional experiences using neurophysiological methods (for more details see the methods section and the past publication [13]). While the measure of physiological parameters such as EDA reflect the emotional arousal associated with music [27, 65], EEG has recently been identified as a promising technique for the investigation of musical reward [2, 12]. During the performances of six candidates conducting musical extracts from the Stabat Mater by F. Poulenc, the participants used a smartphone to report their subjective pleasure according to 4 levels (neutral, low, and high pleasure up to chills; inspired from Salimpoor et al [64]). Additionally, we monitored their emotional arousal by recording Electrodermal Activity (EDA). Thirteen of the 15 participants were included in the analysis, as 2 were excluded because of sensor misrecordings. We also recorded the cerebral activity of twelve of the the participants using mobile EEG (see Figure 1). After each competitor’s performance, the participants were asked to fill out the Aesthetic Experience Scale in Music (AES-M from [57]); a scale that evaluates how people are emotionally “Touched” and “Absorbed” while listening to music as well as occurrences of musical chills. This multimodal hyperscanning approach allowed us to estimate both the cerebral and physiological coupling of pairs of participants involved in the study while they were reporting both similar and different levels of musical pleasure. We used the Total Interdependence (TI) index to estimate inter-individual cerebral coupling [18]. TI is a spectral coherence calculation that quantifies the amount of mutual information between similar electrodes for two people. We also calculated the physiological coupling (Skin Conductance Concordance: *SC Concordance*-[41] throughout the concert for pairs of participants.

**Figure 1:**
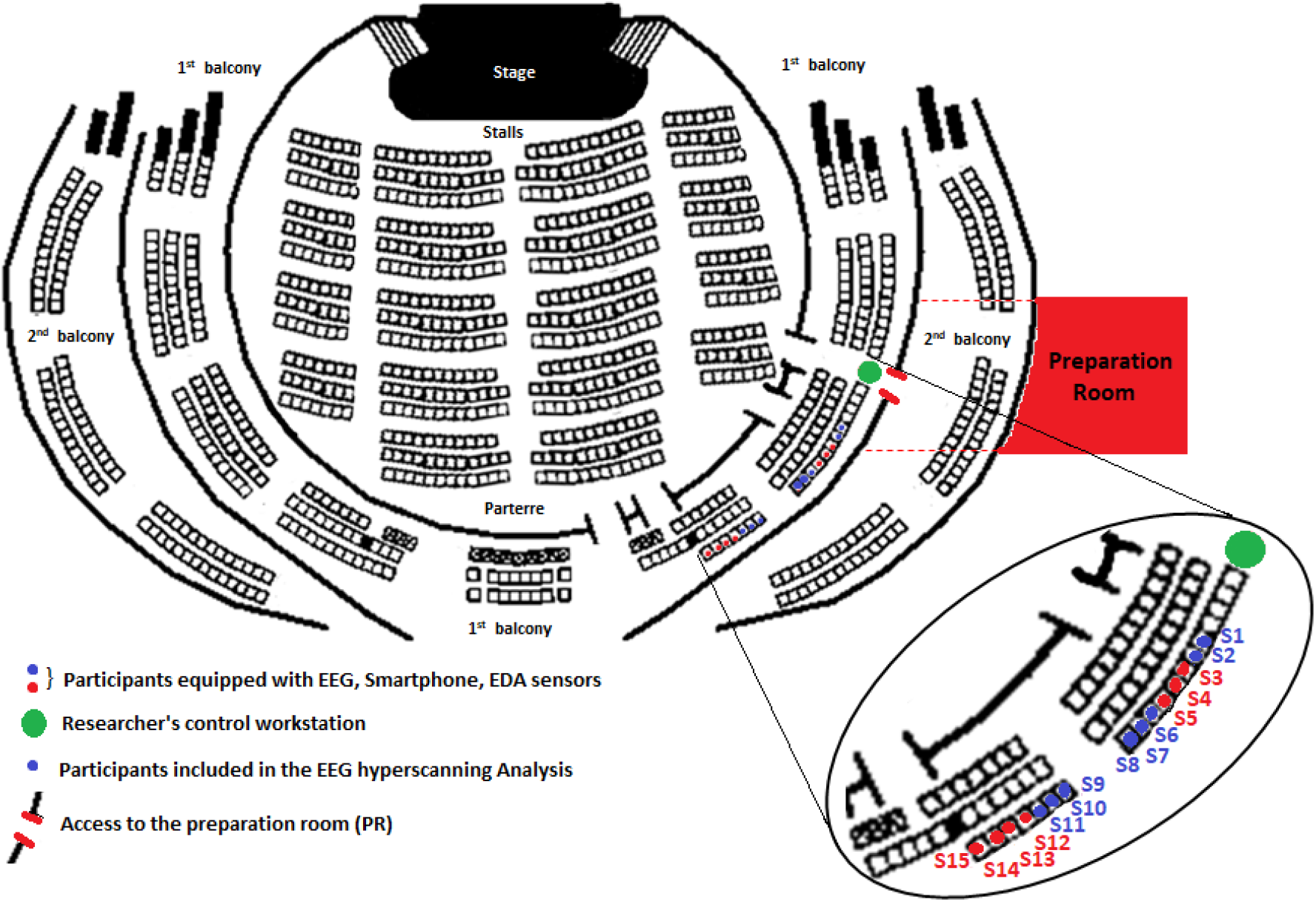
Set up and Positioning of participants in the concert room.

We expected that objective neurophysiological measures of coupling and the behavioral subjective emotional reports would help us to highlight an emotional connection between people during the concert. More precisely, we expected that when reported emotions were similar for several participants, we would observe specific physiological and cerebral coupling between these participants due to activation of similar cerebral structures (e.g. reward system).

## 2 Results

Our analysis of the cerebral coupling shows that the TI was higher when people were listening to music during the concert compared to the baseline period (pause without music), (Wilcoxon signed-rank test: W(66) =1817, p<0.001; for n= 66 pairs, see figure 2-A). Furthermore, the comparison of the TI indexes when participants simultaneously reported either a neutral, low, or high pleasure when the music was played, showed a significant difference (Friedman test; F(2.20)=18, p<1.10^−5^; for n= 21 pairs with enough common periods of signal free from artifacts). The index TI*-neutral* was lower than both the TI*-Low* pleasure and TI*-High* pleasure indexes (respectively p<0.0001 and p<0.001). However, no difference was observed between TI indexes for low and high pleasure (p=0.22) (see Figure 2-B). Not enough chills were reported by pairs of participants simultaneously to calculate a corresponding index for chills.

**Figure 2:**
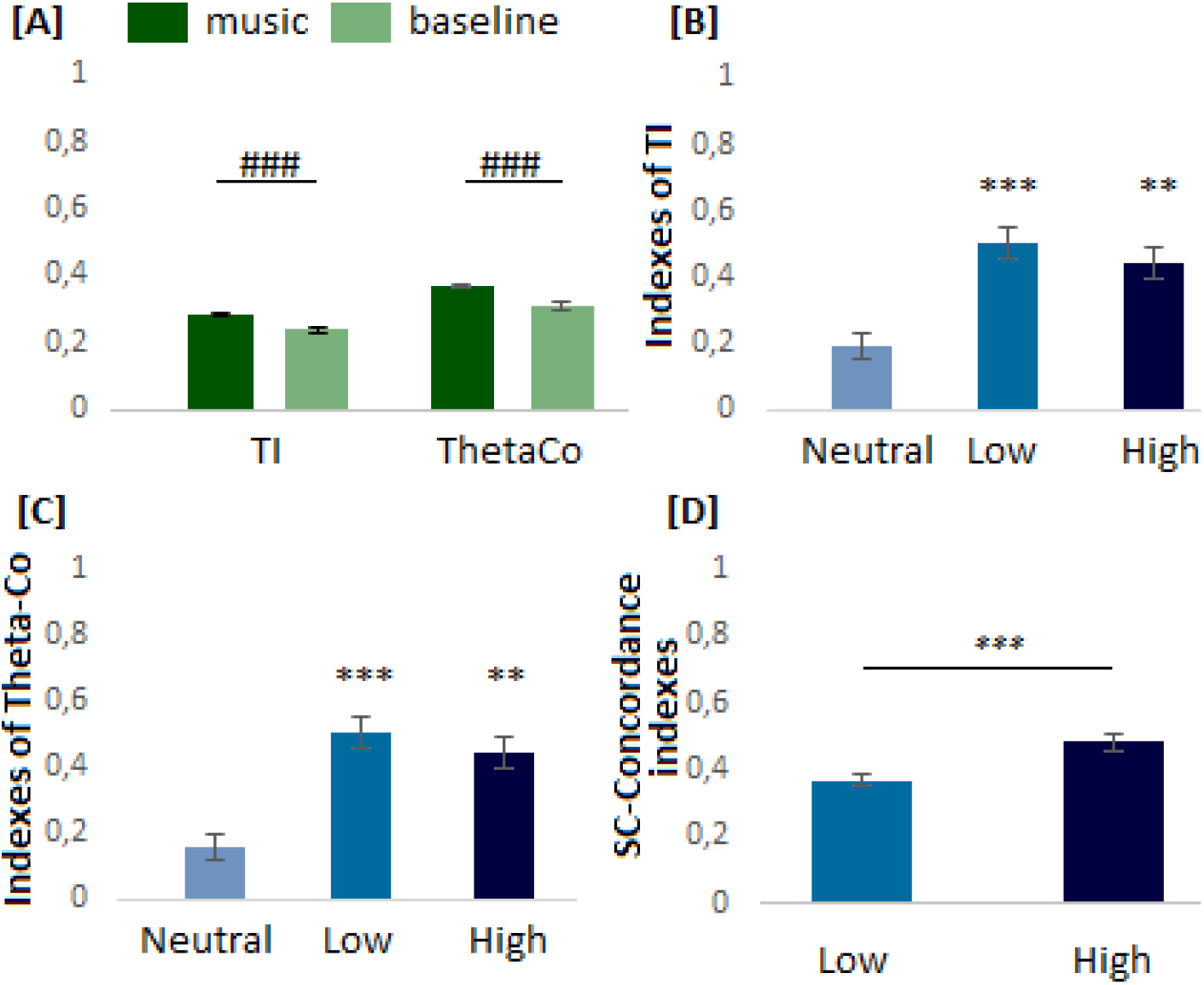
[A] Comparisons between TI and ThetaCo indexes during music vs baseline (# for p<0.05, ### p<1.10^−5^). [B] Comparison of TI indexes and [C] ThetaCo indexes when participants reported simultaneously either a neutral, low, or high pleasure (Significant difference with neutral index; with Bonferonni correction*p<0.016, **p<0.001, ***p<0.0001). [D] Comparisons of Skin Conductance Concordance indexes when participants reported simultaneously a low or a high pleasure (for n=55 pairs, * for p<0.05, ** p<0.01, *** p<0.001).

Since TI is a measure of the cerebral coupling on the whole scalp (AF3, AF4, F4, F3, F7, F8, FC5, FC6, T7, T8, P7, P8, O1, O2) in a wide range of frequencies (1 to 20 Hz), and has been reported to be an estimation of the general engagement with the task [18], we calculated a more specific index of inter-individual emotional coherence. According to recent research in EEG [2, 12], the pleasure derived from music can be estimated on prefrontal and frontotemporal areas in the theta frequency band. On this basis, we calculated an index of theta coherence (ThetaCo) (range of 1 to 4 Hz) on the frontal-prefrontal and temporal electrodes (AF3, AF4, F3, F4, FC5, FC6, F7, F8, T7, T8). In the same way as we observed for the TI indexes, the ThetaCo index while listening to music was higher than the ThetaCo index at baseline (Wilcoxon signed-rank test: W(66) =1864, p<1.10^−5^;, for n= 66 pairs). Additionally, the Friedman Test revealed a significant difference between ThetaCo indexes when participants simultaneously reported their subjective pleasure (F(2.20) = 24.6; p<1.10-5, for n=21 pairs). The ThetaCo index for neutral reports was lower than the ThetaCo indexes for low and high pleasure reports (respectively p<1.10-5 and p<0.001). We observed no significant difference between ThetaCo indexes when participants simultaneously reported low pleasure and when they simultaneously reported high pleasure (p = 0.25) (see Figure 2-C). In addition to the EEG coupling results, we performed a physiological measurement of arousal by recording EDA variations. The Skin Conductance Concordance (SC-Concordance) Index [41] allowed us to estimate the relationship between EDA variations for pairs of participants over time (for more details, see methods section). The SC-Concordance index was significantly higher during music periods than during baseline periods (Wilcoxon signed-rank test: W(77) = 2725; p < 1.10^−8^). In the same way as for EEG, the length of time the participants simultaneously reported either a neutral, low, or high pleasure restricted the number of pairs that could be included in the analysis. We observed that more than 55% of pairs did not report enough simultaneous neutral emotions to be included in the analysis (see SI). Nevertheless, our results show that the SC-Concordance was higher when participants simultaneously reported intense pleasure than when they simultaneously reported low pleasure (Wilcoxon signed-rank test: n=55 pairs; W(54) = 376; p < 0.001; see figure 2-D). Furthermore, the percentage of variation from the baseline when participants reported a high pleasure (mean=1.08 SD=3.43) was higher but not significantly different than when reporting a low pleasure (mean= −3.95; SD=2.28; Wilcoxon signed-rank test: n=13 participants; W(12) = 34; p = 0.45).

To investigate the influence of proximity in emotional sharing between participants, we correlated the TI and ThetaCo indexes with the distance between people (Position Index). We found a negative relationship between the relative position of participants and the cerebral coherence index, but only when people simultaneously reported high pleasure. In other terms, the physically closer the participants, the higher the indexes [Spearman correlations for the index of position vs TI-high: p =0.004, *ρ* = −0.59 - see figure 3-D; index of position vs ThetaCo-high p=0.021, *ρ* = −0.49, see figure S1 in Supplementary Information]. However, we observed no significant relationship between the relative position and the TI and ThetaCo indexes, regardless of emotions reported by participants or when they were simultaneously reporting either low or neutral pleasure [TI-music: p=0.82, *ρ* = −0.05, TI-neutral: p = 0.16, *ρ* = −0.31, TI-low: p = 0.15, *ρ* = −0.32(see Figure 3), ThetaCo-music: p = 0.97, *ρ* = −0.008, ThetaCo-neutral: p = 0.43, *ρ* = −0.17, ThetaCo-low: p = 0.20, *ρ* = −0.28 (See Figure S1 in Supplementary Information)]. We found no relationship between the relative position of the participants and the SC concordance indexes [*Spearman correlation; SC-Concordance-High vs pairs relative position*: p=0.83, *ρ* = −0.029; *SC-Concordance-Low vs relative pairs position*: p = 0.14, *ρ* = 0.19].

Because we observed that the closer the participants, the higher the cerebral coupling for simultaneous intense pleasure reports, we also analyzed the behavioural response variations of pairs of participants (index of Behavioural Synchronization). This index is expressed as a percentage. Higher percentages indicate that participants reported similar behaviors while lower percentages indicate different behaviors (see methods for further details). Our data suggests that the closer the participants were physically, the more they indicated similar levels of emotion simultaneously (Spearman correlation; p = 0.0067, *ρ* = −0.57) see Figure 4-A.

**Figure 3:**
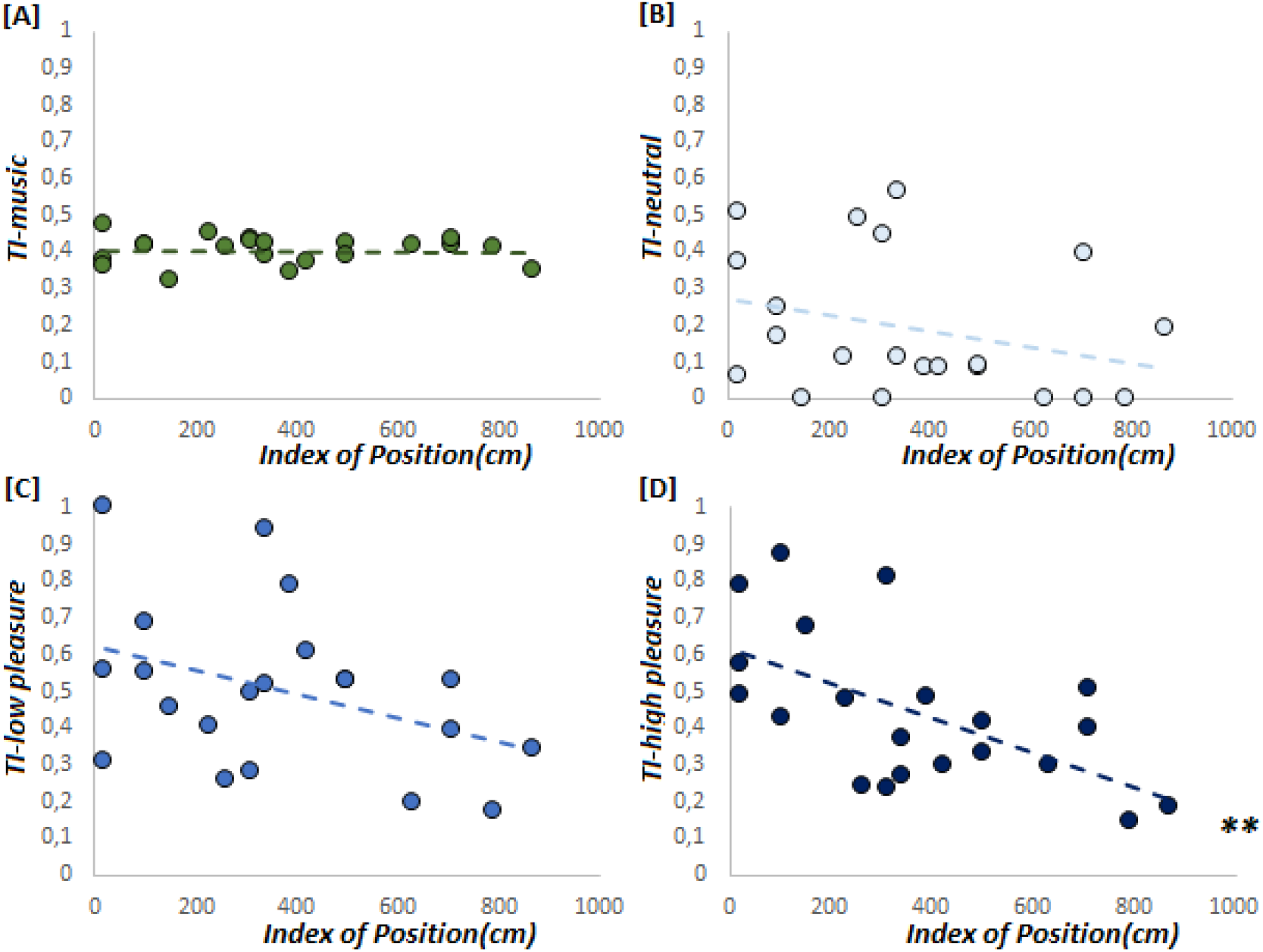
[A] Pearson correlation between the relative position (distance between participants) and indexes of TI; [A] when participants reported different levels of emotion, [B] simultaneous neutral emotions, [C] simultaneous low emotions, and finally [D] simultaneous high emotions (*p<0.05, **p<0.01).

**Figure 4:**
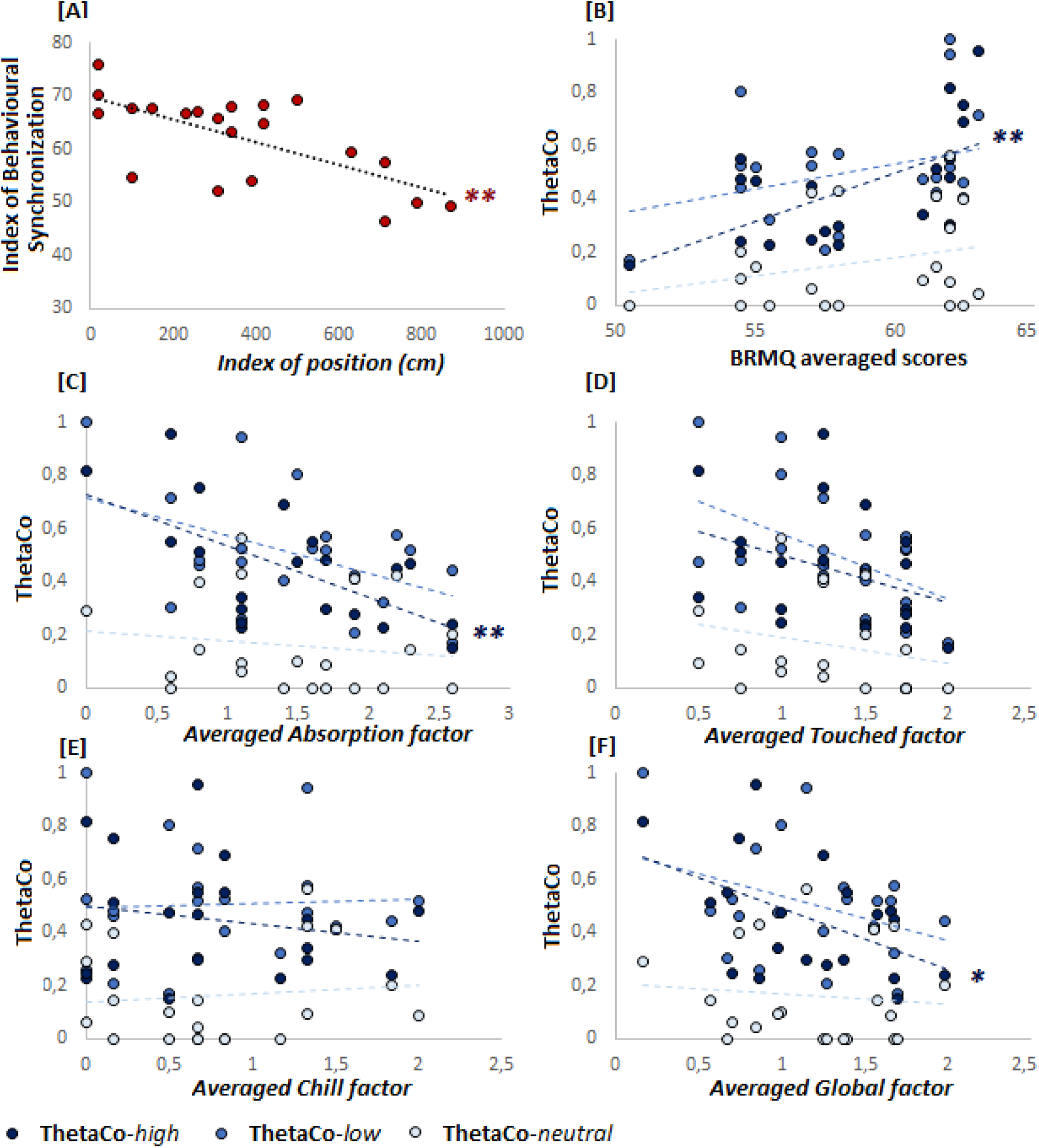
Spearman correlations between [A] the Index of Behavioural Synchronization (IBS) and the index of position, [B] BRMQ pairs average scores and ThetaCo indexes, [C] Absorption factor and ThetaCo indexes, [D] Touched Factor and ThetaCo Indexes, [E] Chill factors and ThetaCo indexes, [F] Global factor and ThetaCo indexes (*p<0.05, **p<0.01, ***p<0.001).

To further understand the link between the participants’ particularities and the coupling results, we correlated the BRMQ averaged scores and the ThetaCo indexes. A positive relationship emerged between the BMRQ averaged scores for pairs and the ThetaCo-high index [p=0.0024, *ρ* = 0.62], suggesting that the higher the BMRQ averaged scores of the pair, the more their cerebral activities were synchronized while they were reporting high pleasure simultaneously. No such relationships were found for the other levels of emotion [BMRQ vs ThetaCo-neutral: p = 0.46, *ρ* = −0.16; BMRQ vs ThetaCo-low: p = 0.53, *ρ* = 0.14, BMRQ vs ThetaCo-music: p = 0.52, *ρ* = −0.14, see Figure 4-B]. Finally, we also correlated the coupling measures and the ratings given with AES-M factors to cross-check neurophysiological results and participants’ subjective emotional experiences. We performed Spearman correlations between the AES-M factors and the ThetaCo indexes. A negative relationship was observed between the Absorption factor and the ThetaCo index when participants simultaneously reported high pleasure [p=0.0033, *ρ* = −0.60, See figure 4-C], and between the Global factor and the ThetaCo-High [p=0.022, *ρ* = −0.49]. For other correlations, no relationship was highlighted (see further results in SI).

## Discussion

We tracked the behavioral, physiological, and cerebral activities of a sample of people attending a concert of liturgical music, with the aim of measuring and understanding how musical emotions are communicated between audience members. We showed that the cerebral activity of physically closer people was more similar when reporting higher degrees of pleasure simultaneously. Since the majority of authors have proposed theoretical approaches to explaining the function of music over decades [68], it seems obvious to address this question by providing new evidence with objective data to test and/or build a case for the most plausible assumptions. The vast majority of the theoretical and empirical research that has studied the function of music has proposed theories or gathered data related to social bonding and social sharing, and descriptives items about the emotional facets of music (see [68] for review). From a broad list of these items, Schäfer et al. [68] constructed 3 major dimensions; “arousal and mood regulation”, “self-awareness” (which includes the emotional aspect of music), and finally “social relatedness”, which is reported to be underestimated. With our approach, we aimed to reconnect the two last dimensions and proposed, like many other authors, that music can be viewed as a social-emotional connector derived from ancestral affective communication [59, 40, 46, 1]. The social context is a catalyst of the musical emotional experience [61]. Furthermore, the involvement of ancient reward circuits as mediators of musical pleasure increases the interest and the complexity of this issue.

Our EEG measures of cerebral coupling mainly suggest that the cerebral activity of people is more similar when they feel intense pleasure during music listening. We expected this result considering that if the same reward structures are activated at the same time for several people enjoying music together, they should simultaneously produce similar cerebral oscillatory activities. Our main interest was in how people can be influenced by the emotions of others. The observation of higher cerebral and behavioral coupling for people that were physically closer suggests a form of emotional connection between participants, or at least a form of inter-individual emotional influence. We proposed to test this finding through measurement of the skin conductance coupling, which is a direct objective witness of parasympathetic activity. As such, we expected that when participants felt an intense pleasure simultaneously, the influence of reward structure activation on the parasympathetic activity would lead to simultaneous skin conductance level increases, thus producing higher physiological coupling. Indeed, the physiological activity varied more similarly for intense than for low pleasure but no relationships were found regarding the relative position of participants. In contrast to the EEG coupling, we explain the absence of a relationship between physiological coupling and position by the slow variations of the skin conductance levels, which are in the range of several seconds.

One possible hypothesis for why people are more “cerebrally” connected when they are physically closer is that the pleasure felt at an individual level can be transmitted between people in the same manner as other basic emotions, i.e. through the mechanism of emotional contagion. Emotional contagion is a mechanism through which an individual can recognize and be synchronized with the emotions of another person via auditory and visual emotional cues (visual contact, vocal emotional cues, and facial expressions, postural, and so on)[6, 31, 67]. It involves mimicry of the perceived emotion, which creates physiological feedback and leads to the subject truly feeling the partner’s emotion [30]. Emotional contagion occurs at a subconscious level involving both the Mirror Neuron System (MNS), which is implicated in the understanding and the imitation of social-emotional behaviour, and the Mentalizing system (MENT), which is implicated in the prediction of the relationship between internal and external states [9, 48, 55, 62].

Firstly, the correlation between the ThetaCo-high and the position index (figure 2-D) seems to show a sort of dropout in the data for distances beyond 2 to 3 meters. This suggests that after 2 or 3 meters when the social contact is limited, the cerebral coherence is lower. Secondly, ThetaCo-high is lower when people are more absorbed in the musical experience. We could argue that the less the participants communicated with peers, the less they transmitted their emotions and the less their neurophysiological rhythms were synchronized. The pleasure felt at an individual level can be transmitted to one or more nearby people not only by simple behavioral mimicry but by emotional expression, which leads to a person being truly emotionally influenced by a peer. This can promote physiological feedback at the individual level, influencing physiological and cerebral rhythms in a similar way. To support this hypothesis, several studies have pointed out that visual contact is a factor of influence of cerebral activity. For example Dikker et al., [18] showed that people who performed a face to face baseline had enhanced cerebral coherence in later activities. Similarly, Leong et al., showed a greater bidirectional influence of infant-adult cerebral activities of direct eye gazes during musical live interactions [38]. Among the vast number of studies that have investigated cerebral coupling using EEG in various domains [73, 21, 75, 18, 38, 33, 10] including the musical context [72, 52, 39, 4, 53], no studies have estimated the emotional cerebral coupling. This lack of evidence in the literature, which could have reinforced our findings, is likely to be mainly due to the difficulty of setting up ecological paradigms and to the lack of objective markers for affective state [15, 14].

Affective or emotional resonance mechanisms could be considered as an alternative explanation for our observations, but these are less obvious interpretations. For now, there are no biological explanations for these mechanisms. The emotional resonance goes beyond the simple emotional recognition leading to a physiological emotional mimesis [49]. It refers to the dynamic processing through which we “move” others’ emotions and “are moved” by the emotions of others (or we are “affecting and being affected”), in social contexts via postural, vocal, and auditory cues [49, 16, 69]. Mühlhoff et al. [49] defined it as the embodiment of social interactions, a physical coupling force, or a “gripping force between” people. The term resonance refers to “what is reverberate”. In physics, it refers to the coupling of two or more dynamic systems that influence each other and are engaged in mutual causal interaction. When system A resonates with system B, each system exerts a mutual oscillatory influence leading to a dynamical coupling force between them. These reciprocal oscillatory activities can enhance the vibratory energy between systems [49]. In social interactions, emotional resonance represents a retro-active loop through which the affect of one individual can reinforce the same affect in another individual [74]. The same data could be interpreted differently since the correlation between position and TI (or ThetaCo) does not appear to have a drop-off after 3 meters, but is rather seen as a gradient. Further experiments should record objective data (e.g; video) to emphasize this inter-individual communication, even in the context of passive listening, which does not normally involve direct continuous interaction between people of the audience.

Music-induced emotions are also related to musical emotional contagion, through which the listener can perceive and even feel the emotions expressed by music and the emotional expression introduced by the composer or the performer [36, 47]. Baltes, and Miu, [5] also argued that when people listen to music in a social context, empathy and emotional contagion might influence the “social facilitation”. We assumed that the surrounding environment of the concert, notably the view of the emotional expression given by the conductors to musicians and the emotional way the musicians are performing, might influence the emotions of participants, but this is not the only possible interpretation of the emotional connection between audience members. A supplementary argument in favor of a more direct emotional communication between participants is that the pairs of participants who were more absorbed in the experience showed lower coherence values. These counter-intuitive results suggest that openness to the surrounding environment helps people to emotionally connect with others. Unfortunately, it is very difficult to estimate the influence of this kind of natural environment on inter-individual emotional communication. It is also difficult to very precisely interpret the effect of the natural variations of the concert (orchestra performance, conductor expressiveness) on the individual emotional experience and the natural inter-individual cerebral coupling. It is well known that familiarity with the musical extracts, which creates anticipation leading to surprises or/and confirmations of expectations is a key factor of pleasure related to reward structure activations [45, 34, 78, 43]. It activates musical memory by bringing back “past encoded musical templates” but also episodic memory through real autobiographical memories in some cases [35]. Among our 15 participants, only 3 reported being familiar with the musical pieces (and were not seated with each other). Considering this low number, we assume that it was not an influencing factor. The affiliation between people was also not a factor of influence since only one pair of participants knew each other (mother/daughter). Furthermore, as previously discussed, the influence of the visual cues on the cerebral coupling and more generally on the cerebral activity is difficult to estimate and represents one of the constraints of a natural environment such as this. A control group outside of the concert room following the concert in a social or individual context could have helped to refine the interpretations. We admit that the constraints for EEG recordings in such natural settings were high and resulted in rejecting a lot of data. For this reason, EEG data included in the analysis was visually inspected to ensure standardized data quality. On the other hand, the identification of noise periods in EDA data is more difficult, which leads to lower data quality.

## Conclusion

In conclusion, our results suggest that people attending a concert together were more connected when they felt emotions or pleasure simultaneously, and that this effect was reinforced when they were physically closer. We suggest that musical emotions felt at an individual level could be influenced by other people through a mechanism of inter-individual emotional contagion (or resonance) and that the mere presence of other people could reinforce this effect. We also discussed our data through the alternative hypothesis of emotional resonance, a physical emotional vibration affecting multiple people in a similar way, considering the strong link between the proximity and the coherence of pairs of participants. These mechanisms are not mutually exclusive and may both have been involved. Further investigation should clarify why people can automatically synchronize with others when similar emotions are experienced and how this effect is constructed on a biological level. The reciprocal interaction between dopaminergic and oxytocinergic pathways in the mesocorticolimbic area during social interactions could be an interesting path for future research.

## Material and Methods

### Participants

The research met the local ethical rules as laid out in French law concerning non-invasive protocols involving healthy participants and was classified as an observational study outside the scope of the Jardé law (Article R1121-1 of the French Law Code of Public Health amended by decree n° 2017-884 of May 2017). It was submitted to the Ethics Committee CPP Est II, which exempted the study from the full ethics review process. Each participant was informed of the observational nature of the study and signed a non-opposition notice designed for the context of observational studies. All people who had bought a ticket for the relevant concert sessions of the International Competition for Young Conductors received an advertising email related to our study. Forty-six people contacted us directly to receive more information. The eligibility criteria for inclusion in the study were to be aged over 18 years old and be right-handed, to be able to wear neurophysiological devices for a significant amount of time, to have a ticket for the relevant sessions of the conductor competition, and to sign the non-opposition notice. A sample of 37 people was included in the study over 4 sessions. The current results concern a sample of 15 participants (12 women) with a mean age of 55.7 years old (SD =18.9, range = 18-78; mean BRMQ scores=56; SD=6.3) recruited specifically for the afternoon semi-final round of the International Competition for Young Conductors on the 18th September 2019. No compensation was given for participation in the study.

### Procedure

During the first meeting, all implications of the study were presented and participants signed the non-opposition notice designed within the framework of observational studies. All were right-handed (Edinburgh Inventory score>50; [58]. Participants filled out the French version of the Barcelona Music Reward Questionnaire (BMRQ) [42, 63]. During this first meeting, all the instructions were explained and a simulation was performed in the laboratory; participants were equipped with mobile EEG headsets and physiological sensors to ensure that the one-size EEG headsets fit the head of every participant for the best possible recordings and to ensure they would be able to tolerate wearing all sensors for long periods during the competition. At the same time, they became familiar with the system of reporting their emotional experience with the smartphone according to four levels of emotion (neutral pleasure, low pleasure, high pleasure, chills) while listening to musical extracts. Based on our experience from the previous tests recording sessions, we asked participants to deactivate the emotional report on the smartphone if they felt displeasure while listening to the music, in order to not introduce a measurement bias within the neutral level. This simulation was necessary to explain to the participants what is required and what is not acceptable for neurophysiological recordings. Participants were asked to stay calm and to adopt a comfortable position, to avoid head or eye movements, and to avoid any motor movements not necessary for realtime behavioral reports. During the concert, 15 participants were equipped with both the smartphone and the physiological sensors (Shimmer Sensing GSR 3 + unit for HRV and EDA recordings) and 12 of these participants were additionally equipped with a Emotiv Epoc + EEG headset. They were prepared in the adjacent preparation room and were then comfortably installed in the first balcony of the concert hall on a single row of seats (see Figure 1). This specific positioning allowed us to calculate a position index, which represents the distance between people (see Figure 1). The participants then watched 6 conductors in turn conducting a full symphonic orchestra and choir, performing extracts from Stabat Mater of Francis Poulenc (*I.Stabat Mater Dolorosa, II.Cujus animam gementem, IV.Quae moerebat, VII.Eja Mater, XII.Quado Corpus*). Each performance lasted approximately 25 minutes including repetitions and instructions to musicians.

### Analysis

#### Behavioral report & Questionnaires

we assessed the behavioral synchronization between participants by calculating an index that measures the variations of behavioral reports over time. The index of Behavioural Synchronization (IBS) is expressed as a percentage and takes the value of 100% when participants reported the same level of pleasure. The value was fixed at 50% when participants reported levels of pleasure that differed by 1 point (eg; high pleasure versus low pleasure). Finally, the value was fixed at 25% when participants reported levels of emotion that differed by two points and 0% for differences of more than 2 points. Continuous reports during the musical periods allowed us to calculate the index with precision to the nearest second for the whole performance. The participants also completed the Barcelona Music Reward Questionnaire [42, 63], which evaluates sensitivity to musical reward. They also completed the Aesthetic Experience Scale in Music questionnaire extracted from Nusbaum and Silvia [57] after each contestant’s performance (as in [13]). For the correlational analysis, we averaged the questionnaires scores’ of each participant of a pair.

#### EEG

EEG data were low-pass and high-pass filtered (Butterworth) between 1 and 30 Hz, with a notch filter fixed to 50 Hz using Cartool software. Regression of ocular artifacts was performed according to the method from Dikker et al., [18] (more information in Supplementary Information (SI)). The full data was cut into periods of one second, each of which was visually inspected to reject period not free from artefacts. We then calculated cerebral coupling using two indexes; an index of Total Interdependence (TI, calculated over 1 to 20 Hz on 14 electrodes; AF3, AF4, F3, F4, FC5, FC6, F7, F8, T7, T8, P7, P8, O1, O2), which is related to global engagement in the task, and the index of Theta Coherence (ThetaCo; calculated over 1 to 4 Hz on frontal, prefrontal, and temporal electrodes; AF3, AF4, F4, F4, FC5, FC6, F7, F8, T7, T8), which we linked with emotional sharing. For the inter-individual cerebral coupling analysis, the statistical unit becomes the pairs of participants (66 pairs for the 12 participants). We calculated the ThetaCo index specifically on frontal/prefrontal and temporal areas, since several studies have highlighted that theta oscillations over these areas are involved in musical pleasure processing [2, 12, 54, 66].

The calculation of these indexes was based on Dikker et al., and Wen et al., methods[18, 76], they estimate the stability of the relationship between two signals in the frequency domain. TI (formula given in Figure S2 Supplementary Information (SI)) measures “the amount of mutual information between two systems”, with 0 when signals are independent and 1 when there is a strong linear relationship between signals [76]. The calculation is applied between the same electrodes for two subjects and the mean of each electrode index value is calculated to provide a global index. The index values were normalized with a min-max transformation across all pairs of subjects. Each index was calculated using a minimum of 60 acceptable common periods for both participants of the pair and was calculated on musical periods greater than 40 seconds. The ThetaCo index was calculated using the same method. Different conditions were estimated for each index; first, we calculated a “baseline” condition; periods when no music was played by the orchestra during each conductor’s performance (the participants were told to avoid movements, discussion, or other parasitic activities even during this time). Then we calculated “music” conditions when the music was being played (without taking into account the emotion reported). We calculated these conditions for TI and ThetaCo with all the 66 available pairs. Then 3 emotional conditions were calculated for both TI and ThetaCo; the high pleasure condition (or High) when participants simultaneously reported high emotion, and the low and neutral pleasure conditions when they reported either low or no specific musical pleasure simultaneously. Among the twelve participants, the minimum of 60 common periods for cerebral coupling index calculation restricted our analysis to only 21 of the 66 pairs. For other pairs, the participants did not report enough similar emotion or did not have enough common periods that were free from artifacts.

#### EDA

First the Skin Conductance (SC) data was filtered using a low-pass filter fixed to 15.6 Hz and median filter smoothing. The EDA’s measures of coupling were applied to 13 participants. Two participants were not included in the EDA analysis because of data loss due to sensor misrecordings. To estimate the coupling of the Skin Conductance activity between pairs of participants, we used the method from MAci et al.,[41]. We used a Pearson correlation to estimate the SC-Concordance. We made moving windows of 5 seconds (with 1-second overlap) and calculated Pearson correlations on 15-second periods (corresponding to 15 slope averages with 1-second increments) between SC values from pairs of participants. Next, we calculated a global index of SC coupling for each pair by estimating the ratio of the sum of the positive correlations divided by the sum of the absolute value of negative correlations for the whole experiment. We applied a natural logarithmic transformation to each pair’s indexes because of the skew related to ratios. Finally, we applied a min-max transformation. An index value close to 0 indicates that a low physiological concordance was observed between the two participants, while a value close to one indicates a high physiological concordance. We only included in the analysis pairs of participants for which a minimum of 2 minutes of coupling could be calculated. This analysis requires that participants report similar emotions during a sufficient time. Then, as for the EEG measures of coupling, we calculated coupling indexes when participants simultaneously reported a neutral, low, or high pleasure while listening to music.

## Acknowledgments

The authors thank the director of the International Festival of Besançon Franche-Comté; Jean Michel Mathé and his team, for opening the doors of the International Competition for Young Conductors with enthusiasm and without restrictions. The authors also thank Tanawat Chansophonkul, Lisa Michelant, Charline Compagne, Coralie Joucla, Yvan Peterschmitt, Pierre-Edouard Billot, Nathan Galmes, Julian Sanchez, and Alpha Ntumba for their kind support for experiment planning and data acquisition. The authors also thank Jennifer Dobson for proofreading the article, and Gregory Tio and Sebastien Michea for their help in this research.

The study was carried out using funds belonging the CIC-IT 1431 and the platform Neuraxess, with material and the human support from the Integrative and Clinical Neuroscience Laboratory of Besançon. The study also benefited from a doctoral grant from the French Ministry of Higher Education and Research.

## Contributions

T.C., D.G., and L.P., designed research; T.C., D.G., A.C., and L.P., performed research; T.C., D.G., and A.C. analyzed data; T.C., D.G., A.C., T.M., E.H., and L.P., wrote or/and revised the paper.

## Author Declaration’s

The authors declare no conflict of interest.

