## supplementary material for "Your pleasure is mine; when people share a musical emotional experience during a live music performance in a concert hall"

March 2021

#### Supplementary Information

### Results

#### Correlational analysis full report

In the results section, we highlighted some correlations between Theta-Co indexes and factors of the AES-M scale. The complete statistical report is presented hereafter: Absorption factor vs ThetaCo-neutral: p = 0.55, $\rho$ = -0.13; Absorption factor vs ThetaCo-low: p = 0.17, $\rho$ = -0.31; Absorption factor vs ThetaCo-music : p = 0.79, $\rho$ = 0.059; Touched factor vs ThetaCo-neutral: p = 0.07, $\rho$ = -0.39; Touched factor vs ThetaCo-low: p= 0.07, $\rho$ = -0.39; Touched factor vs ThetaCo-high: p= 0.05, $\rho$ = -0.42; Touched factor vs ThetaCo-music: p= 0.62, $\rho$ = -0.11; Chill factor vs ThetaCo-neutral: p = 0.83, $\rho$ = 0.04; Chill factor vs ThetaCo-low: p= 0.88, $\rho$ = 0.033, Chill factor vs ThetaCo-high: p= 0.77, $\rho$ = -0.06; Chill factor vs ThetaCo-music: p= 0.81, $\rho$ = -0.05; Global factor vs ThetaCo-neutral: p=0.59, $\rho$ = -0.12; Global factor vs ThetaCo-low: p=0.28, $\rho$ = -0.24; Global factor vs ThetaCo-music p=0.89, $\rho$ = -0.02.

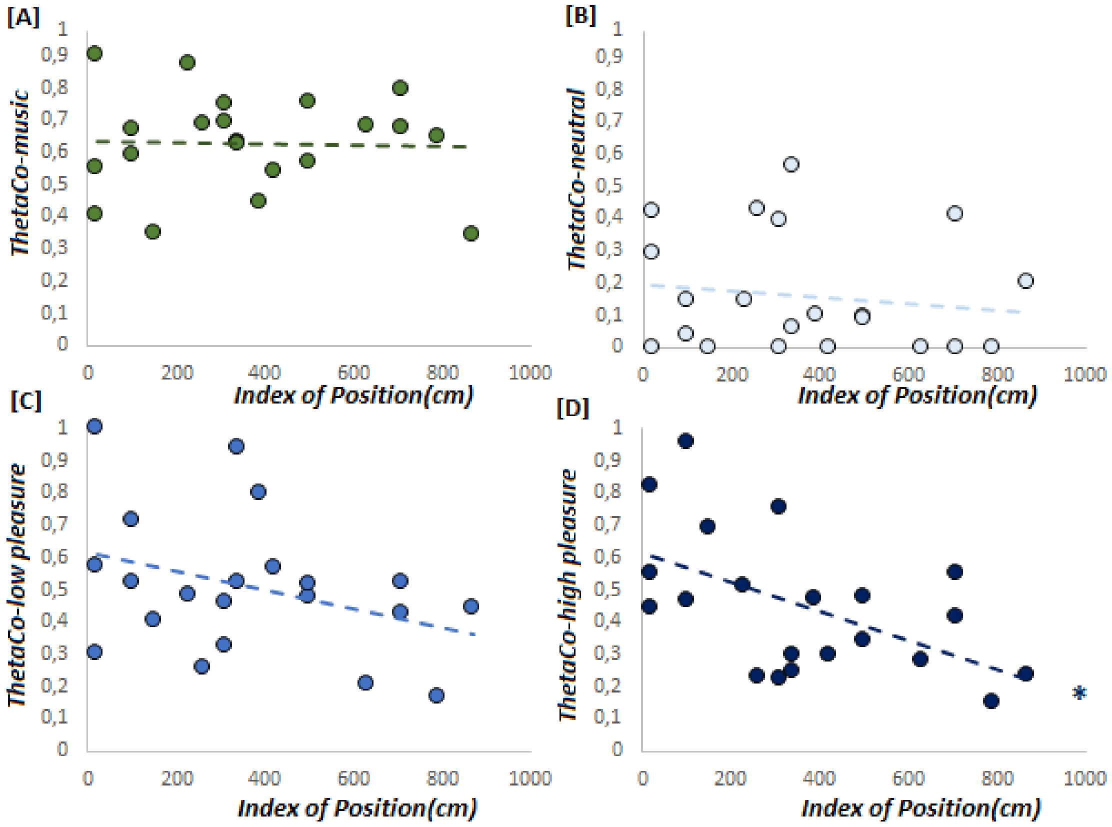


Figure 1: [A] [A] Spearman correlation between the relative position (distance between participants) and ThetaCo indexes; [A] when participants reported different levels of emotion, [B] simultaneous neutral emotions, [C] simultaneous low emotions, and finally [D] simultaneous high emotions (*p<0.05, **p<0.01).

### Method

#### 1.Acquisition system and device synchronization

The acquisition system was set up by Manaty Cloud Team. Octopus is a software that allows users to record events in a database from multiple distant sources with a precise central time reference.
It is opensource, developed in Java, and is designed to run as a cluster on machines that might not be on the same network.
Each node of the cluster comprises:
- A standalone program (octopus daemon) that manages the cluster interaction for registration of nodes and time synchronization, serves the web client, and gather data from multiple sources.
- A local database where events are stored
- A set of connectors that handle the connection to external data sources and collect data.
- A web client (JS frontend) that allows a human user to monitor the cluster and the event recording sessions.

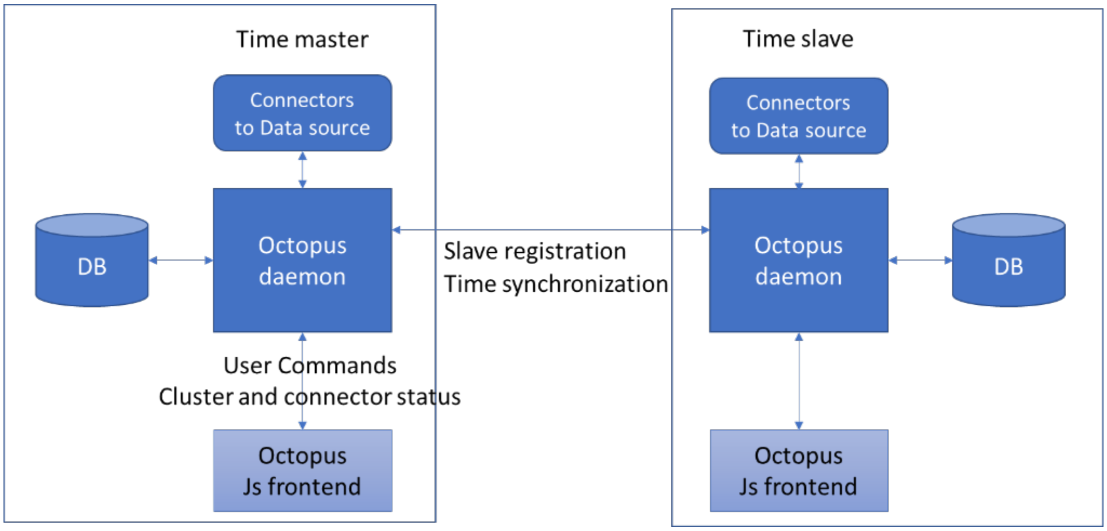


Connectors have been implemented for :
- Emotiv Epoc headsets
- Dedicated Android emotion recording app
- User manual input from the web client

Each node records the event it receives or collects in a local database. It is only after the recording session is over that the events are exported and merged in a single storage. It is therefore of tremendous importance that all the nodes in the cluster are aware of the time difference between them. A time difference computation process between each slave and the master, consisting of a batch of ping-pong requests and response over a TCP socket is performed regularly during the recording session. Each slave is, therefore, able to send events in the database with its local time and the time difference with the master. The master is also responsible for triggering the start and end of the recording session on each node and gathering the status and metrics of all the connectors of each node so they can be monitored on the web client of the master node.

Regarding the synchronization of EDA data, the software Consensys Pro provided by Shimmer Sensing allows synchronization of the internal clocks of each device (GSR unit) to the time of a reference PC via the Shimmer Base. Thereafter, data is recorded on the SD card of the device and resynchronized during the importation on the reference PC. This allowed us to synchronize the EDA, behavioral and EEG data based on the same timestamp. Unfortunately SD card recorings did not give access to a real-time data plot.

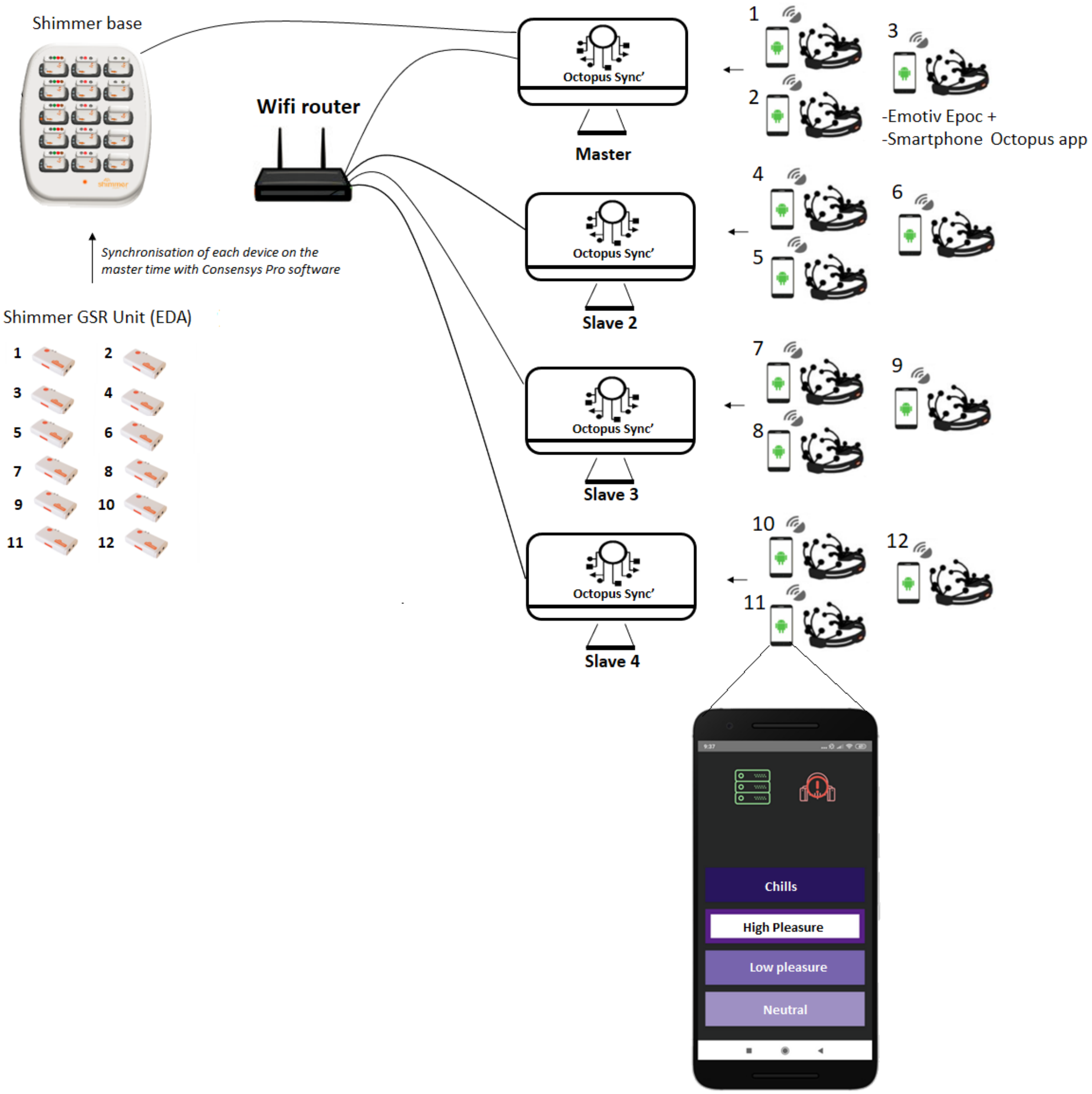


Figure 2: Architecture of the synchronized acquisition system for behavioral, electroencephalographic, and physiological data.

2. EEG data selection for measures of coupling

The Bluetooth connection of EEG devices was sometimes lost during the recordings. That results in several data loss. The table S1 present the percentage of data loss by participants for each candidate

#### 3.EEG Ocular artifacts regression

We led independent research (already published **?**), to ensure that the mobile EEG headset Emotiv Epoc + used in this study could provide relevant information about musical emotion and musical pleasure mechanisms in the frequency domain.

Since eye blink could influence the inter-individual coherence values when they are rhythmically produced by an external source such as the rhythmic movement of the conductor, the regression of ocular artifacts is a major concern. We performed a Principal Component Analysis (PCA) according to the method from Dikker et al., **?**, which used the same Emotiv Epoc + wireless devices. An estimation of the amount of eye blink artifacts is made based on the electrodes close to the eyes. F7 and F8 are parallel to muscles responsible for horizontal movements of the eyes, and thus became a good predictor of horizontal eye movements. Similarly, AF4 & F8 and AF3 & F7 predict vertical eye movements. The PCA allowed us to regress the collinearity between EEG data and an eye movement predictor and then to build a linear regression model.

**EEG$_{PCA}$ = (model constant) + (slope coefficient x eye movement predictor) + EEGresidual$_{PCA}$**

EEGresidual$_{PCA}$ is not related to eye movement. The EEG$_{PCA}$ model predicted the EEGresidual$_{PCA}$ value, which is the main component that cannot be estimated by the eye movement predictor is calculated. We were then able to back-project the main component of the EEGresidualPCA into the EEG data of the electrode domain by applying the inverse of the coefficient of the main component.

**EEG = (principal component coefficient)$^{-1}$ x EEGresidual$_{PCA}$**
This procedure was performed on all EEG sections for all participants.

#### 3. EEG data Analysis

After the regression of the ocular artifacts, data were re-synchronized and down-sampled to 125 Hz based on the timestamps. All the signal was cut into periods of one second. Each period was visually inspected to reject every epoch not free from artifacts. Then all periods in common were used to calculate the indexes both when the music was played and during periods of pause (Indexes of TI and ThetaCo-music and pauses).

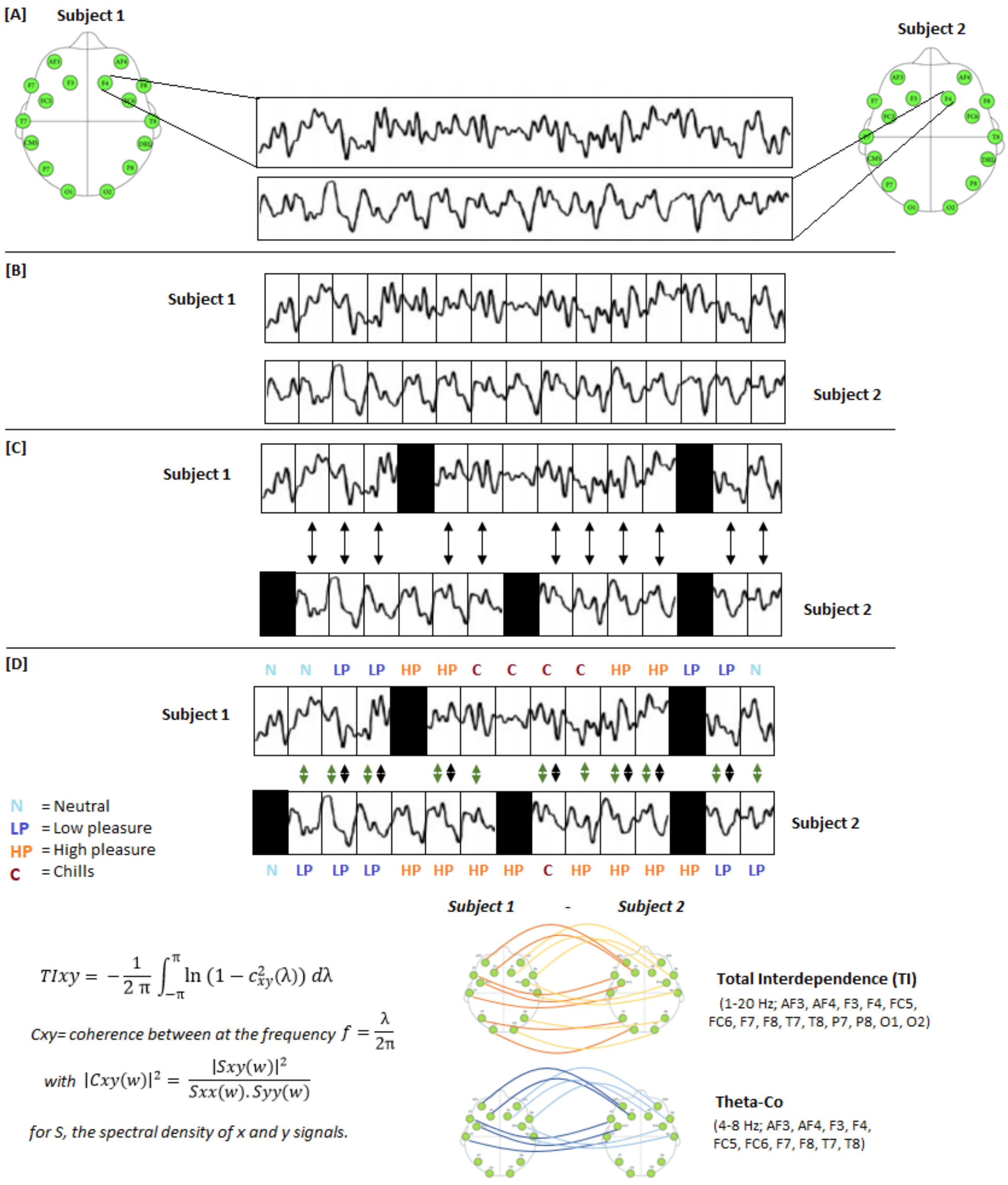


Figure 3: Steps of the EEG data Analysis; after the eye movement regression, [A] the EEG signal is extracted for each similar electrode for each pair of participants and synchronized in the range of the millisecond. [B] The signals are split into one-second epochs. [C] Epochs not free of artifacts are rejected and the analysis is performed on the remaining common usable epochs. [D] Coherence analyses of TI and ThetaCo are performed both on all common periods and on periods for which the participants indicated the same levels of pleasure (black arrows).
